# Missed Opportunities of Chlamydia and Gonorrhea Detection when not Using Extra Genital Screening Among Males

**DOI:** 10.1101/428706

**Authors:** Chelsea Schafer, Belinda Prado, Nora Barin, Leslie Gama

## Abstract

**Background:** Through extra genital screening methods, Health Departments and community clinics can increase detection of sexually transmitted diseases (STDs) through the application of urethral testing. The Long Beach Department of Health and Human Services (LBDHHS) works on preventing cases *Chlamydia trachomatis* (CT) and *Neisseria gonorrhoeae* (GC) infection from being undiagnosed, by providing extra genital screening.

**Methods:** Retrospective medical review of 1,569 patient health records, who received CT/GC testing, based on at least one visit to the LBDHHS, STD Clinic, from 2012 to 2015. All male patients ages 18 years or older with positive CT/GC results (n=242) for urethral, rectal, and pharyngeal sites; regardless of their sexual behaviors, were included in the study. Females, those under the age of 18, and patients who tested negative for all three anatomical sites were excluded (n=1,327).

**Results:** At time of collection, study participants had a mean age of 37 years. Reported ethnicity indicated 56% Caucasian, 21% Hispanic or Latino, 9% Asian or Pacific Islander, 7% Other, 5% Black, and 2% More than one race. The use of extra genital screening detected 15% (n=242) of the 1,569 patients tested positive for at least one type of CT/GC infection. These findings demonstrated that if urine were the only specimen collected, then over 29.7% of CT and 46.8% of GC cases would have been missed.

**Conclusions:** Testing of all three anatomical sites should continue to be performed for CT/GC detection. Cases of CT/GC are underreported if performing urethral screening alone. These testing measures may reduce the potential for missing a diagnosis, mitigating transmission, as well as prevention and control.

**Short Summary:** The number of *Chlamydia trachomatis* (CT) and *Neisseria gonorrhoeae* (GC) infections missed were determined using extra genital screening procedures among male patients from the Long Beach Department of Health and Human Services (LBDHHS), STD/Family Planning Clinic. This method tests urethral, pharyngeal, and rectal sites for CT and GC infection. In this retrospective medical review of 1,569 patient files, of which 242 tested positive, the proportion of male cases that would have remained undetected for CT/GC if using urethral screening only were detected.

## Introduction

Sexually transmitted diseases (STDs) have been an ongoing public health concern as communicable disease rates have steadily increased. From 2012-2016, CT cases increased by 51% (530 to 801 per 100,000), while rates of GC have increased by 214% (98 to 309 per 100,000). In 2016, Long Beach, California experienced the second highest rate of Chlamydia (CT) and third highest rate of Gonorrhea (GC) reported for the State of California. The City of Long Beach faces one of the highest morbidity rates of CT and GC in the state ^(1)^. Infection can occur during intercourse, when there is contact with the bacterium *Chlamydia trachomatis* and/or *Neisseria gonorrhoeae* ^(2)^. Once infected, individuals may experience pain during urination, discharge from penis, and swollen testes. Infection can be detected in the genitals, rectum, and throat. However, it is possible to show no signs or symptoms of infection, being asymptomatic ^(3–4)^. Testing of these STDs is critical to determine a person’s infection status and can be detected in the genitals, rectum, and throat ^(2–4)^. Based on 2017 STD & HIV Screening Recommendations from the CDC, annual CT/GC screenings should be performed among all sexually active males, regardless of condom use ^(5)^. Screening is recommended every 3-6 months if the individual has multiple or anonymous sex partners. Despite CDC recommendations, there continues to be a lack of performing extra genital screening. According to the CDC, one in three clinicians have not heard of extra genital screening ^(6)^. Previous research identifies most patients do not report any type of sexual behavior to their clinicians, leading to underreporting of infections being contracted ^(6–8)^. This could be due to patient’s status being asymptomatic. Men who have sex with men (MSM) are disproportionately affected for contracting STDs. CT/GC can increase their risk for co-infections such as human immunodeficiency virus (HIV) ^(9)^. Young males are at greater risk for infection if residing in areas with a high prevalence of CT ^(5, 10)^. Extra genital screening, also known as triple-site testing, has been the method of choice for CT/GC detection, as it examines pharyngeal, rectal, and urethral sites with first-void urine samples using nucleic acid amplification tests (NAATs). It is considered the most sensitive in regard to specificity for cell culture and recognition of bacterial sample assays ^(8, 11)^. There is currently no standard recommendation on screenings for GC among heterosexual males ^(5)^.

In 2017, the California Department of Public Health reported rates of CT to be 552.2 per 100,000 population, with Los Angeles having a rate of 560 per 100,000 population. That same year, rates of GC were 190.3 per 100,000 population in California ^(3, 12)^. Communicable disease rates have been on the rise, calling for improved methods for early detection and STD testing. Screening programs for these harmful infections are made widely available to reduce the burden of disease in the population. Extra genital screening tests patients in their genital, urethral, and pharyngeal sites. Although urine is most widely used to test for CT/GC infection, it is still recommended to test all three sites as an individual’s sexual practices can place them at greater risk for infection, depending on the type of sex they are having ^(13, 15)^. Information on the underlying disparities regarding prevalence and incidence of disease is necessary for bridging the gap in services. These issues could be related to differences in cultural backgrounds and risky sexual behaviors such as lack of condom use or high number of sex partners ^(13–14)^.

In a previous study conducted by Kent, Chaw et al. ^(8)^ at a health clinic in San Francisco, researchers noticed that if patients were only screened at urethral sites, 53% of CT and 64% of GC infections would have been missed. In the same study, among MSM population, more than 70% of CT infections would have been missed if they only tested for GC infection. This would have been based on a false assumption that among those who tested positive for GC infection, that patients were also tested for CT infection ^(8)^. Other researchers have reported similar findings stating that if a clinic does not perform extra screening measures, then roughly 70% of CT/GC cases, would have gone unreported ^(16–17)^.

Due to high rates of STDs in the City of Long Beach, there is increased need for improved detection, patient education, and surveillance of CT/GC among residents. Extra genital screening of CT/GC detects more cases of infections, when using nucleic acid amplification tests (NAATs), enhancing sensitivity and specificity ^(13, 18)^. STD testing is performed opportunistically, regardless if patients show any signs or symptoms, since many of the cases seen are asymptomatic ^(3)^. Examining high-risk populations, based on their sexual behaviors and infection status, is most cost effective and has demonstrated a reduction in the incidence of CT/GC infections ^(3)^. The purpose of this study was to determine the number of CT/GC cases that would have been missed and left untreated if not performing extra genital screening among male patients who were tested at the LBDHHS, STD Clinic. Patient and provider education are needed to increase awareness, while reducing stigma associated among those seeking STD testing and treatment. The transmission and susceptibility of CT/GC infection is reported to be enhanced among patients with HIV ^(19)^. Performing urethral screening alone creates the opportunity for missed infections in other anatomical sites. Health Departments and STD Clinics should continue to perform extra genital screening of all three anatomical sites as there is limited data at the national level, underestimation of disease prevalence, and risk for HIV ^(16–17)^. STD screening guidelines should continue to be followed to reduce incidence of new cases.

## Methods

Researchers conducted a retrospective medical review of 242 cases who identified as male and tested CT/GC positive after extra genital screening. A total of 1,569 patients were tested for CT/GC at the LBDHHS, STD Clinic during a 4-year period, from 2012 to 2015. Study participants were chosen based on whether they received STD testing during at least one visit to the on-site clinic at LBDHHS. Females, those under the age of 18, and patients who tested negative for all three anatomical sites were excluded (1,327 controls). All male patients ages 18 years or older with positive CT/GC results (242 cases) for any of the three sites, including urethral, rectal, and pharyngeal; regardless of their sexual behaviors, were included in the study. All health records were collected during patient intake with no attempt to re-contact. Written consent was provided at the time of the physical exam. Patients’ medical charts were pulled from NextGen v5.6 to assess data on STD test results and demographics. The LBDHHS lab performed internal validation of nucleic acid amplification testing (NAAT) to screen extra genital sites, as well as urine testing to test for CT/GC positivity. In particular, sensitivity and specificity of STD testing is reported to be significantly enhanced by application of molecular techniques with NAATs ^(18)^. All patients were evaluated in a clinical setting and diagnosed in accordance to California Department of Public Health (CDPH) guidelines, with serological confirmation ^(10)^. Test positivity was defined as the proportion of males who had positive extra genital and/or urogenital CT/GC test (n=242) ^(15, 20, 23)^.

To determine the amount of missed CT/GC infections, analysis was done using Pearson’s Chi-Square test for association to generate a cross-tabulation and frequency tables using SPSS Statistics 24 ^(15, 20, 23)^. The number of cases who tested positive per disease at site of infection were assessed, including those positive for both pharyngeal and rectal, yet negative urethral sample ^(15, 20, 23)^. Prior exposure was assessed using recent sexual contact to determine symptoms of CT and/or GC. Sexual behaviors were recorded into categorical variables, with patients grouped as either men who have sex with men (MSM), men who have sex with women (MSW), or men who have sex with men and women (MSMW). For operationalization measurement during health record review, variables were chosen if having exhaustive and mutually exclusive categories. To increase the likelihood of the successful achievement of the study objective, application of sample size selection using threshold probability of a (two-tailed) α test =.05, β =.20, and r=.20, indicated a minimum sample size of 194 cases for statistical significance ^(21)^.

## Results

At time of collection, age of patients ranged from 15-75, with a mean age of 37 years. 52.5 % (n=127) identified as “*Not Hispanic or Latino(a)*,” 42.6% (n=103) “*Hispanic or Latino(a)*,” and 5.0% (n=12) “*Unknown or Not Reported*” for ethnicity. Patients reported race indicated, 56% (n=135) White, 21% (n=52) African American or Black, 9% (n=21) Asian or Pacific Islander, 7.4% (n=18) Other, 5% (n=11) Hispanic or Latino, and 2% (n=5) More than one race. Results indicated that an estimated 29.7% (n=50) of CT and 46.7% (n=73) of GC infections would have been missed if only performing urethral testing. There were 5.9% (n=10) pharyngeal, 20.8% (n=35) rectal, and 2.9% (n=5) pharyngeal-rectal cases that showed positive for CT and negative by urine (Table 1 - Appendix A). In addition, 16.0% (n=25) of pharyngeal, 16.7% (n=26) rectal, and 14.1% (n=22) pharyngeal-rectal cases tested positive for GC and negative by urine (Table 2 - Appendix B). To determine if there was a difference in the amount of cases reported from 2012 to 2016, a percent change calculation was performed. This demonstrated an increase of CT rates by 51% (530 to 801 per 100,000) and GC rates by 214% (98 to 309 per 100,000).

## Discussion

These findings provide data on the incidence and prevalence of CT/GC cases in The City of Long Beach. Results from these analyses show the need for implementing extra genital screening, in addition to urine testing, among male patients in clinical settings to prevent cases from going undiagnosed. The use of extra genital screening with NAATs may improve detection of asymptomatic cases or those that would otherwise be undiagnosed if a patient does not report their sexual health history. Testing of all three anatomical sites should be performed routinely at health clinics to detect for infection early since patients can be asymptomatic. Fewer cases of CT/GC are to be reported among sexually active males. It is recommended to continue these practices across clinical settings to control the spread of infection and ensure the appropriate treatment is being offered. Due to lack of data at the national level, and provider education on extra genital screening, there is underestimation of CT/GC prevalence ^(6, 16)^. About one in three clinicians have not heard of extra genital screening, which has resulted in the lack of performing these testing procedures ^(6)^.

Study limitations included selection bias through the use of convenience sampling methodology. Distinguishing temporal precedence was determined to be a consequence of using retrospective health records for assessment. Early detection and treatment of these bacterial infections can help in the prevention and control of other STIs, such as HIV and HPV. The Health Department is one of the most well-suited entities for having a surveillance program, as they can reach the most vulnerable populations through targeted interventions. However, there is still a lack of performing extra genital screenings for CT/GC, even if indicated. It is recommended by the Centers for Disease Control and Prevention (CDC) that annual screenings be done based on those at greatest risk for infection ^(19)^.

Evaluation of CT/GC is important for ensuring that the appropriate medical tests are being done, patients receive adequate treatment, and a reduction STD rates. This study contributes to the field of epidemiology by providing information on CT/GC prevalence among males. Extra genital screening needs increased support as a recommended STD testing procedure. This is especially true among MSM, due to sexual practices that put them at increased risk for infection such as anal intercourse. Despite national organizational recommendations, more support is still needed if wanting to include this STD testing and screening method for other populations such as females. Surveillance data might not be well representative of CT testing and reported rates. Findings from researchers Kustec, et al. ^(22)^ indicate underreporting of CT cases among females to the National Institute of Public Health (NIPH) ^(22)^. Health programs should bring increased awareness, while removing stigma associated among those seeking STD testing and treatment. This can be done by providing routine extra genital screening at Health Department Clinics for prevention and control. Testing of all three anatomical sites can reduce opportunities for missed infections, as fewer cases of CT/GC are more likely to be reported if urethral testing only. Further research is still necessary to determine the amount of cases who receive treatment and follow through on their CT/GC infection status.

## Appendix - Tables

## Appendix A - Table 1

**Table 1.** Chlamydia (CT) by Extra Genital Site among Male patients, Long Beach, 2012-2015 (n=242)

|  |  |
| --- | --- |
| **Extra Genital Screening Test Results** | **Number of Cases** |
| *Pharyngeal (negative), Rectal (negative), and Urethral (negative) | 118 |
| Pharyngeal (positive) and Urethral (negative) | 10 |
| Pharyngeal (positive), Rectal (positive), and Urethral (negative) | 5 |
| Rectal (positive) and Urethral (negative) | 35 |
| Urethral (positive) | 74 |
| **Total (n):** | **242** |
| **Patient reporting Positive Test site, negative by Urethral:**<br>**PH (10) + PH Rectal (5) + Rectal (35)** | **50** |
| **Total number of cases (242) – Urethral Positive (74)** | **168** |
| **% Missed CT** | **29.7% (50/168)** |

This table includes all cases who tested positive for a single site, those who tested positive for two sites (Rectal & Pharyngeal) and those who tested negative for all three sites.
- Those who were negative by Urethral, and negative/not done for Pharyngeal and/or Rectal test.

## Appendix B - Table 2

**Table 2.**
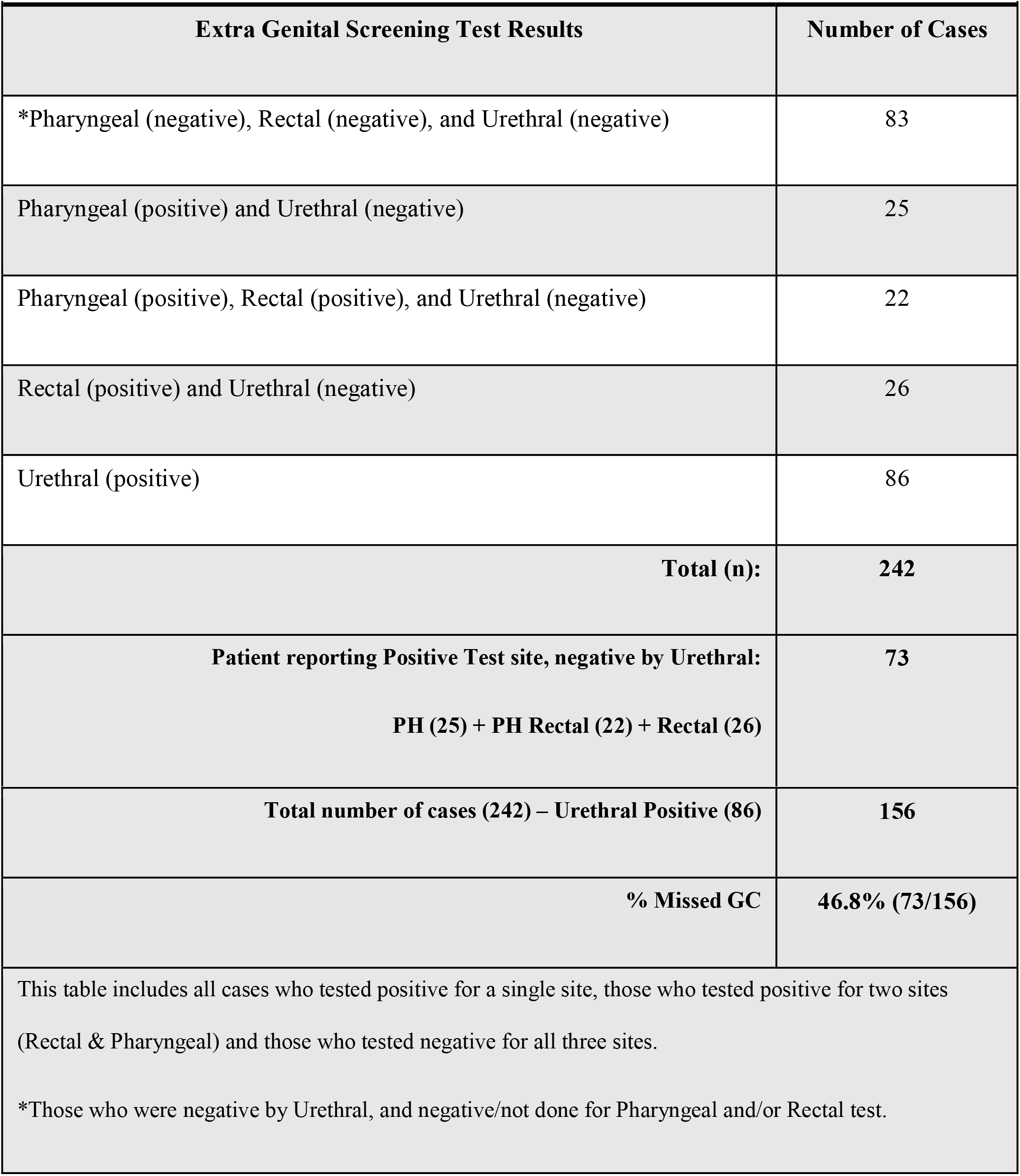
Gonorrhea (GC) by Extra Genital Site among Male patients, Long Beach, 2012-2015 (n=242)

## Notes

#### Summary of Updates

The number of Chlamydia trachomatis (CT) and Neisseria gonorrhoeae (GC) infections missed were determined using extra genital screening procedures among male patients from the Long Beach Department of Health and Human Services (LBDHHS), STD/Family Planning Clinic. This method tests urethral, pharyngeal, and rectal sites for CT and GC infection. In this retrospective medical review of 1,569 patient files, of which 242 tested positive, we determined the proportion of cases that would have remained undetected if only using urethral screening.

