## Supplementary material for "Missed Opportunities of Chlamydia and Gonorrhea Detection when not Using Extra Genital Screening Among Males": Cover Letter (from another version of this preprint: version 2, 2019-01-27)

Statement of Disclosure:

There is no known conflict of interest that exists for purposes of this study.

Sources of Support:

We would like to acknowledge the California Department of Public Health, Centers for Disease Control and Prevention, and the City of Long Beach with special thanks to the Health and Human Services Department.

### Cover Letter

Researchers examined the effectiveness of using extra genital screening (e.g. urethral, pharyngeal, and rectal) procedures for *Chlamydia trachomatis* (CT) and *Neisseria gonorrhoeae* (GC) detection, by assessing the number of cases that would have been missed if only using urethral screening. Researchers conducted a retrospective review examining 1,571 MSM patient health records. These patients received extra genital screening for CT/GC at the Long Beach Health Department, STD Clinic, between 2012 and 2015. Among 1,571 cases, 242 (15%) patients tested CT and/or GC positive for at least one anatomical site, including urine, pharyngeal, and/or rectal. Our findings demonstrated that if urine was the only specimen collected, then 29.7% of CT and 46.8% of GC cases would have been missed. This study demonstrates the increasing rates of CT/GC and high percentage of cases that would have gone undiagnosed without extra genital screening.

The principal investigator, Chelsea A. Schafer, has a background in public health with a Bachelor of Science in Health Education and Epidemiology Intern for the City of Long Beach, Department of Health and Human Services. She maintains her Certificate as a Certified Health Education Specialist (CHES) and is program planner to the American Public Health Association (APHA), Veterinary Public Health Special Interest Group.

Co-investigator, Belinda Prado, MPH, is the HIV/STD Surveillance Coordinator at City of Long Beach, Health and Human Services Department. Prior to the position, she served as Communicable Disease Epidemiologist for LBHHD.

Co-investigator, Nora Barin, MPH, worked as a Community Health Worker, specializing in Epidemiology for the City of Long Beach, Health and Human Services Department before becoming an Epidemiologist. She also holds her Bachelor's Degree in Public Health from the University of California, Irvine.

Leslie Gama, is a Health Education intern for the for the City of Long Beach, Health and Human Services Department. Leslie Gama graduated from California State University, Long Beach in 2018, where she received her Bachelor of Science degree in Public Health. It is her hope that the information presented serves as a tool to highlight the importance of extra genital testing among men.
